# Dissolved organic carbon (DOC) is essential to balance the metabolic demands of North-Atlantic deep-sea sponges

**DOI:** 10.1101/2020.09.21.305086

**Authors:** Martijn C. Bart, Benjamin Mueller, Titus Rombouts, Clea van de Ven, Gabrielle J. Tompkins, Ronald Osinga, Corina P.D. Brussaard, Barry MacDonald, Anja Engel, Hans Tore Rapp, Jasper M. de Goeij

## Abstract

Sponges are ubiquitous components of various deep-sea habitats, including cold water coral reefs and deep-sea sponge grounds. Despite being surrounded by oligotrophic waters, these ecosystems are known to be hotspots of biodiversity and carbon cycling. To assess the role of sponges in the carbon cycling of deep-sea ecosystems, we studied the energy budgets of six dominant deep-sea sponges (the hexactinellid species *Vazella pourtalesi*, and demosponge species *Geodia barretti, Geodia atlantica, Craniella zetlandica, Hymedesmia paupertas* and *Acantheurypon spinispinosum)* in an ex situ aquarium setup. Additionally, we determined morphological metrics for all species (volume, dry weight (DW), wet weight (WW), carbon (C) content, and ash-free dry weight (AFDW)) and provide species-specific conversion factors. Oxygen (O_2_) removal rates averaged 3.3 ± 2.8 *µ*mol O_2_ DW_sponge_ h^−1^ (all values mean ± SD), live particulate (bacterial and phytoplankton) organic carbon (LPOC) removal rates averaged 0.30 ± 0.39 *µ*mol C DW_sponge_ h^−1^ and dissolved organic carbon (DOC) removal rates averaged 18.70 ± 25.02 *µ*mol C DW_sponge_ h^−1^. Carbon mass balances were calculated for four species (*V. pourtalesi, G. barretti, G. atlantica* and *H. paupertas*) and revealed that the sponges acquired 1.3–6.6 times the amount of carbon needed to sustain their minimal respiratory demands. These results indicate that irrespective of taxonomic class, growth form, and abundance of microbial symbionts, DOC is responsible for over 90 % of the total net organic carbon removal of deep-sea sponges and allows them to sustain in otherwise food-limited environments on the ocean floor.

## Introduction

The oceanic seafloor constitutes by far the largest part of Earth’s surface area, covering an area of 361 million km^2^, of which over 90% is found at water depths greater than 150 m (Costello et al. 2010; Ramirez-Llodra et al. 2010). However, still only a minute fraction of the deep-sea surface has been properly mapped (Mayer et al. 2018), let alone characterized in terms of biodiversity and ecology. Nevertheless, since 1848, 28 new habitats have been discovered in the deep-sea (Ramirez-Llodra et al. 2010). In the past few decades, sponges have been revealed to be ubiquitous inhabitants of many of these habitats, generally in water depths between, but not restricted to, 200–2,000 m (reviewed by Hogg et al. 2010). On the northern Atlantic continental shelf, sponges abundantly inhabit deep-sea coral reefs, form large mono-specific sponge grounds, and create sponge reefs by depositing thick spicule mats (i.e. layers of skeletal needles derived from dead and damaged sponges) (Thomson 1873; Klitgaard & Tendal 2004; Buhl-Mortensen et al. 2010; Beazley et al. 2015). In some areas, sponges can comprise up to 98 % of the total benthic biomass and sponge abundance amounts up to 24 individuals per m^2^ (OSPAR Commission 2010). Deep-sea sponges are found to fulfill important ecological roles by providing habitat complexity and substrate to both mobile and sessile fauna (Klitgaard, 1995; Beazley et al. 2013; Hawkes et al. 2019).

Moreover, the first estimations on respiration and organic carbon (C) uptake of deep-sea sponges (e.g., Pile & Young 2006; Yahel et al. 2007; Kahn et al. 2015) suggest that they play a crucial role in benthic-pelagic coupling.

However, due to technical restrictions inherent to deep-sea work (e.g., costly ship-based expeditions, sampling under extreme conditions), data on the ecology and physiology of deep-sea sponges is still scarce, and mostly based on specimens caught with dredges and trawls. In recent years, the increased use of remotely operated vehicles (ROVs) has provided more opportunities to do measurements at the seafloor, and to bring up specimens from depth for laboratory experiments. The few available studies on deep-sea sponge physiology consists of a mix of in situ and ex situ studies using different direct (taking in- and out-flow water samples (Pile & Young 2006; Yahel et al. 2007; Leys et al. 2018)) and indirect (using flume experiments (Witte et al. 1997), or incubation chambers (Kutti et al. 2013, 2015; Rix et al. 2016)) methodologies. Still, metabolic rates of deep-sea sponges are only available for a limited number of species, often incomplete, and not reflecting the diversity and wide array of morphological traits found in sponges in deep-sea habitats.

Deep-sea sponges mainly belong to two classes: demosponges (Demospongiae) and glass sponges (Hexactinellidae) (Lancaster et al. 2014). Demosponges come in a wide variety of shapes and sizes, ranging from mm-thin encrusting sheets to m-wide barrels, and consist of layers of specialized cells (Simpson 1984). They occur in freshwater and marine ecosystems and their skeleton can consist of siliceous, calcium carbonate, or collagenous components (Müller et al. 2006; Ehrlich et al. 2010; Bart et al. 2019). Hexactinellids are exclusively marine, tubular, cup-, or vase-shaped, and predominantly inhabit deep-sea habitats (Schulze 1887; Mackie & Singla 1983; Leys 2007). In contrast to demosponges, their cellular structure is principally composed of massive multinucleate syncytia and their skeleton always consists of silica spicules (Bidder 1929; Mackie & Singla 1983; Leys 1999; Müller et al. 2006).

Depending on the quantity and composition of associated microbes in their tissues, sponges can be further classified as having either low microbial abundances (LMA) or high microbial abundances (HMA) (Hentschel et al. 2003; Weisz et al. 2008). LMA sponges contain microbial abundances and sizes comparable to ambient seawater (∼ 0.5–1 ⨯ 10^6^ cells mL^−1^), while HMA sponges can contain up to four orders of magnitude more (and generally much larger) microbes (Vacelet & Donadey 1977; Reiswig 1981; Hentschel et al. 2003). These symbionts are involved in various processes, such as C and nitrogen (N) metabolism, synthesis of vitamins, chemical defense and horizontal gene transfer (reviewed by Pita et al. 2018).

Sponges, including deep-sea species, are well-established filter feeders, efficiently capturing and processing nano- and picoplankton (reviewed by Maldonado et al. 2012). More recently, it has been shown that many shallow-water sponges primarily rely on dissolved organic matter (DOM) as food source (reviewed by de Goeij et al. 2017). DOM, often measured in the form dissolved organic carbon (DOC), is the largest potential food source in the oceans (Hansell et al. 2009). Yet, direct evidence of DOM uptake by deep-sea sponges is still not available at present. For some species DOM uptake has been suggested (Leys et al. 2018), for others it was not found (Yahel et al. 2007; Kahn et al. 2015). However, these studies did not directly measure DOC, but derived the dissolved organic carbon fraction from the total organic carbon fraction. Direct DOC measurements are challenging, as they are performed almost within detection limits of current analytical systems. Therefore, an important question is: can deep-sea sponges utilize DOM as a food source?

Both body shape and microbial abundance are suggested to affect the capability of sponges to utilize dissolved food sources. For example, it is hypothesized that the high surface-to-volume ratio of flat, encrusting sponges is advantageous for the uptake of DOM compared to lower surface-to-volume ratio of erect, massive (e.g., ball, cylinder) growth forms (Abelson et al. 1993; de Goeij et al. 2017). Higher DOM uptake is also predicted for HMA sponges in comparison with LMA sponges, as microbes are considered to play an essential role in the processing of DOM (Reiswig 1974; Freeman & Thacker 2011; Maldonado et al. 2012; Hoer et al. 2018). However, this distinction is not always clear, as the diet of some LMA sponges also consists mainly of DOM (e.g., de Goeij et al. 2008; Mueller et al. 2014), particularly when they do not have massive growth forms (reviewed by de Goeij et al. 2017).

To quantify the metabolic- and carbon removal rates of deep-sea sponges, three aspects need to be investigated. Firstly, a wider variety of sponge biodiversity (e.g., different taxonomic classes, morphological shapes, and abundances of associated microbes) needs to be included in physiological studies. Despite the high sponge diversity present in the deep-sea, most studies currently available focused on a single sponge class or species. Secondly, the potential role of DOM in the diet of deep-sea sponges needs to be assessed. Thirdly, sponge metabolic rates are currently normalized to a variety of metrics (e.g., dry weight, wet weight, volume, or organic carbon content), which affects general ecological interpretations and that makes it difficult to compare results between studies. Therefore, the use of different standardization metrics should be addressed.

In this study, we investigated the oxygen and organic carbon removal rates of six dominant North-Atlantic deep-sea sponges with different morphological traits (three massive HMA demosponges, two LMA encrusting demosponges, and one massive LMA hexactinellid) from two different habitat types (mono-specific sponge grounds and multi-specific sponge assemblages associated with cold-water coral reefs), and composed carbon budgets. Specifically, we studied the removal rates of live particulate organic carbon (LPOC; i.e. bacterio- and phytoplankton), DOC, and dissolved oxygen (O_2_) of the investigated species, using ex situ incubation experiments. We further determined different morphological metrics for the six targeted species (volume, DW, WW, carbon content, and AFDW) and provide species-specific conversion factors.

## Materials & Methods

### Study areas, sponge collection and maintenance

We investigated the following dominant North-Atlantic deep-sea sponge species (Fig. 1 and supplementary Table S1): *Vazella pourtalesi* (Hexactinellida; LMA; massive vase), *Geodia barretti* (Demospongiae; HMA; massive, globular), *Geodia atlantica* (Demospongiae; HMA; massive, bowl), *Craniella zetlandica* (Demospongiae; HMA; massive, globular), *Hymedesmia paupertas* (Demospongiae; LMA; encrusting, sheet) and *Acantheurypon spinispinosum* (Demospongiae; LMA; encrusting, sheet). Sponge specimens were collected by ROV during four research cruises in 2016, 2017 (two cruises), and 2018 (Fig. 2). *V. pourtalesi* specimens were collected in August, 2016, attached to their rocky substrate at ∼300 m depth, during the Hudson cruise 2016-019 (Kenchington et al. 2017) at the Emerald Basin on the Scotian Shelf, Canada (44°19’8.73” N 62°36’18.49” W). Sponges were kept in the dark in a 1000-L holding tank and transported without air exposure to the Bedford Institute of Oceanography, Dartmouth, Nova Scotia, Canada. In the lab, sponges were kept in the dark in a 1000-L flow-through holding tank, through which sand-filtered seawater from the adjacent Bedford Basin was continuously pumped at 7 L h^−1^. A chiller was used to maintain water temperature at 8 °C. *C. zetlandica* specimens were collected during the Kristine Bonnevie cruise 2017610 (April 2017) at 60°42’12.5” N 4°39’09.9” E in the province of Hordaland, Norway, and kept in 14-L onboard flow-through aquaria with seawater pumped through at 120 L h^−1^. Temperature was maintained at 8 °C. *G. atlantica* and *A. spinispinosum* specimens were collected attached to their rocky substrate during the G.O. Sars cruise 2017110 (August 2017) at the Sula reef (64°42’25.2” N 7°59’24.0” E) of the Northern Norwegian coast at depths of 250–400 m. *G. barretti* specimens were collected during the same cruise at the Barents Sea (70°47’20.8” N 18°03’47.2” E) at a depth of 272 m. The latter three sponge species were kept on board the research vessel in the dark in 20-L flow-through tanks in a climate room at 6 °C. North-Atlantic seawater was pumped in from a depth of 6 m at 30 L h^−1^. *H. paupertas* specimens were collected attached to rocky substrate during the G.O. Sars cruise 2018108(cruise code, August 2018) in the Barents Sea at 70°47’13.9” N 18°03’23.8” E. These sponges were kept on board the research vessel under similar conditions as during the previous year. Ex situ experiments with *G. atlantica, A. spinispinosum* and *H. paupertas* specimens were performed on board the ship. *G. barretti* and *C. zetlandica* specimens were transported without exposing them to air to the laboratory facilities at the University of Bergen, Norway, where the experiments took place. In Bergen, sponges were kept in a dark climate room (8 °C) in multiple 20-L flow-through aquarium systems. Each holding tank contained a maximum of five sponges. Flow originated from unfiltered water pumped from 200 m depth from the outer fjord near Bergen at ∼ 50 L h^−1^ with a temperature ranging from 6–8 °C. All sponges and substrate were cleared from epibionts prior to incubations.

**Figure 1.**
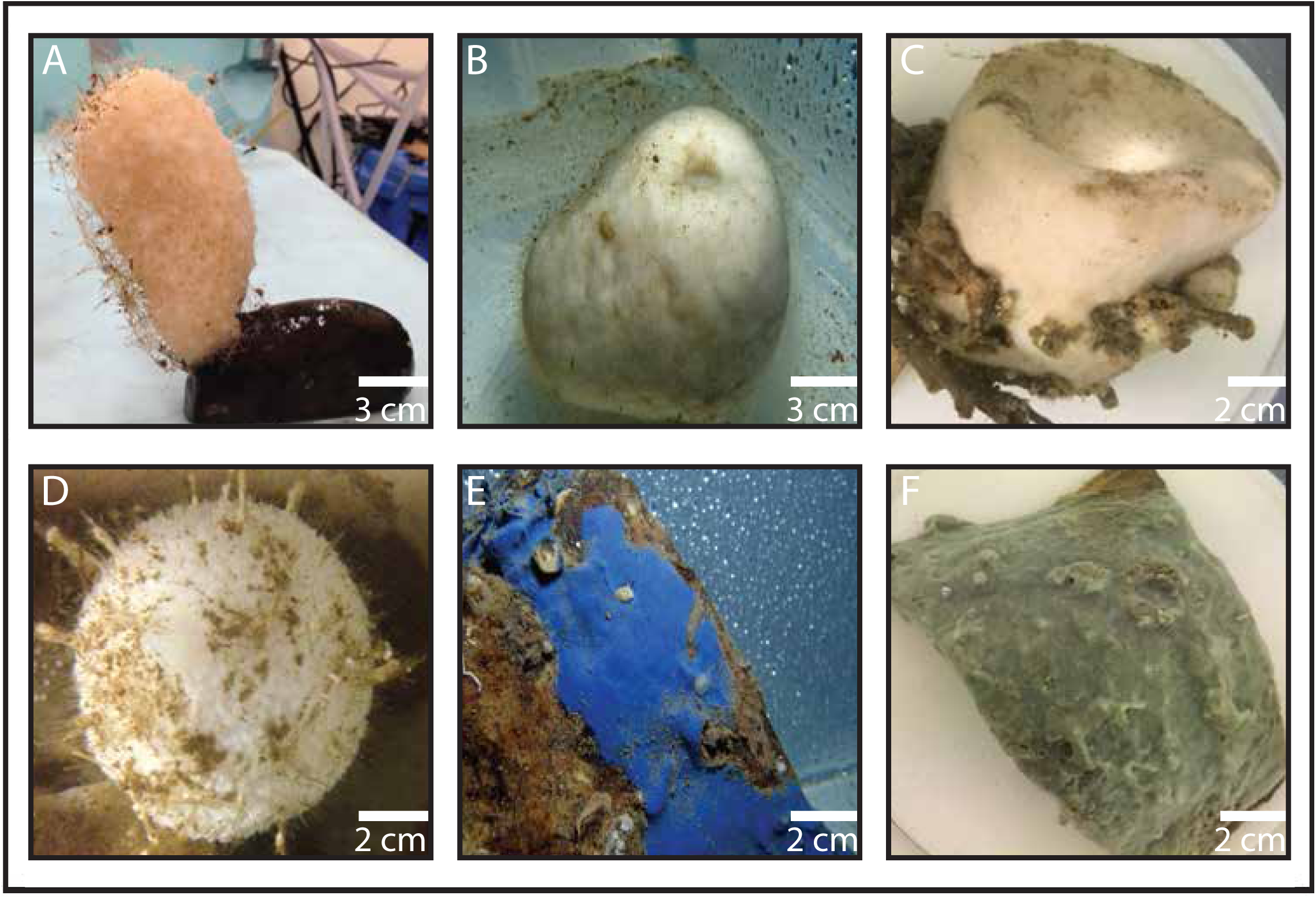
Photographs of six dominant North-Atlantic deep-sea sponge species used in the study. (**A**) *Vazella pourtalesi* (**B**) *Geodia barretti* (**C**) *Geodia atlantica* (**D**) *Craniella zetlandica* (courtesy of Erik Wurz) (**E**) *Hymedesmia paupertas* (**F**) *Acantheurypon spinospinosum*.

**Figure 2.**
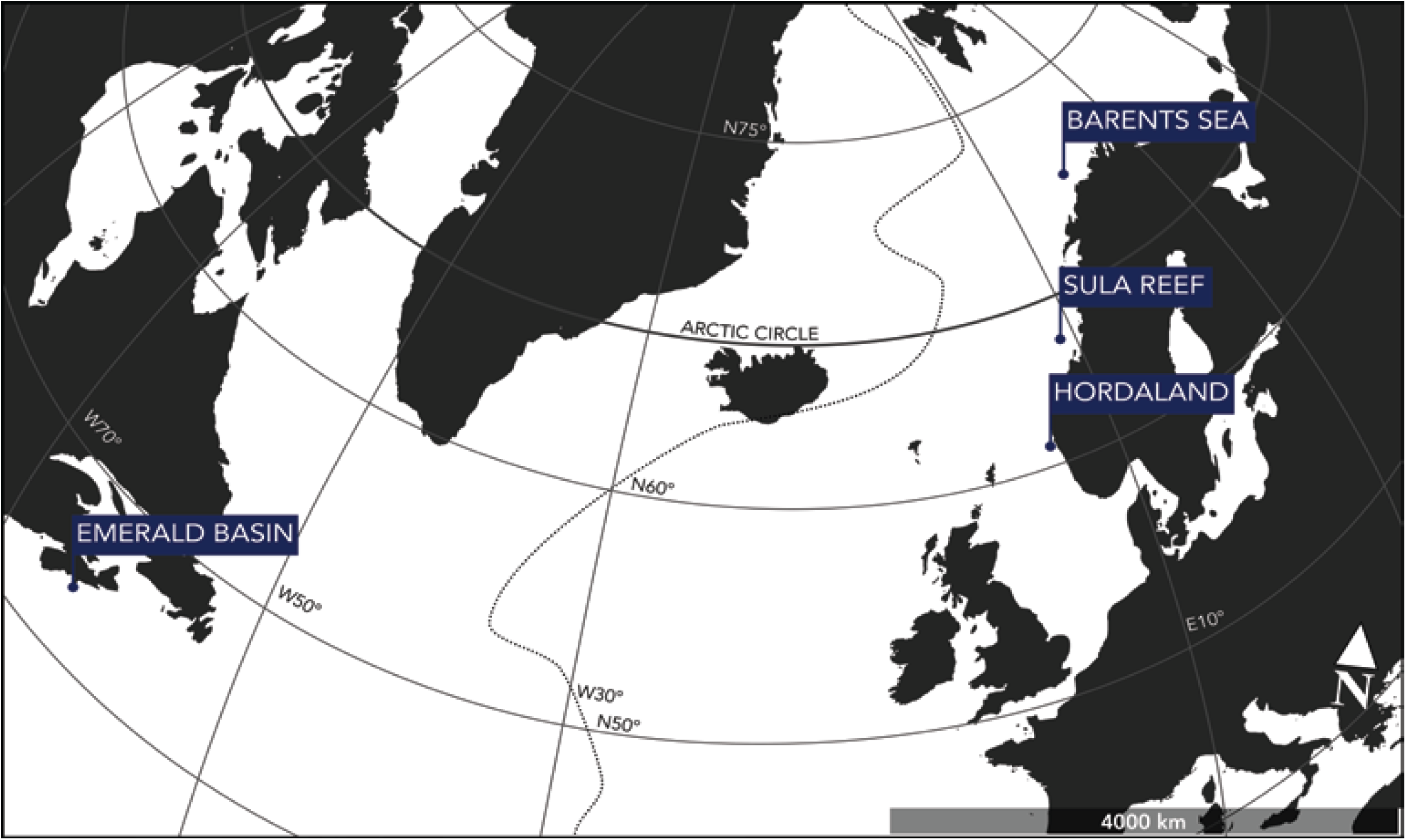
Study area. Sponge specimens were collected at different locations in the North Atlantic during 4 different research cruises in 2016 (Emerald Basin), 2017 (Hordaland, Sula Reef, Barents Sea) and 2018 (Barents Sea). The dotted line represents the Mid-Atlantic Ridge (MAR).

### Incubations, sample treatment, and analysis

All sponges were allowed to acclimatize for a minimum of 1 week prior to the incubation experiments. Individual sponges were enclosed in flow chambers (either 2, 3, or 6 L depending on sponge biomass) with magnetic stirring devices to ensure proper mixing (de Goeij et al. 2013). Chambers were acid-washed (0.4 mol L^−1^ HCl) prior to the incubations and kept in a water bath to maintain a constant seawater temperature during the incubations (6–9 °C depending on the incubation). Chambers were closed without trapping air in the system. The length of every individual incubation was determined during test incubations based on sponge size and oxygen consumption (ideally timed to about > 10 % to < 40 % [O_2_] decrease, and ranged from 2–8 h. At set time intervals depending on the incubation length (t_sample_ = 0, 30, 60, 90 120, 180, 240, 360 or 480 min), 85–100 mL water samples were taken with acid-washed 100-mL polycarbonate syringes. Sample water volume was replaced aquarium water (drawn in by suction to maintain volume and eliminate air exposure), and calculations were adjusted to take these replacements into account during the analysis. Samples were subdivided to analyze the concentrations of DOC and abundances of bacterio- and phytoplankton. For 2 species, *C. zetlandica* and *H. paupertas*, no DOC samples were analyzed, due to logistical issues.

Dissolved oxygen concentrations (O_2_) were continuously measured during the incubations with OXY-4 mini optical oxygen sensors (*PreSens*, Germany). Sensors do not consume oxygen and due to their small dimensions (Ø 2 mm), flow and mass-transport inside the chambers are not disturbed. O_2_ concentrations were recorded every 15 s (*OXY-4-v2_30FB software*).

Prior to DOC sampling, syringes, glassware and pipette tips were rinsed three times with acid (8 mL 0.4 mol L^−1^ HCl), three times with milli-Q (80 mL), and twice with sample water (10 mL). 20 mL of sample water was filtered (< 20 kPa Hg suction pressure) over pre-combusted (4 h at 450 °C) GF/F glass microfiber (∼ 0.7 *µ*m pore-size) filter and collected in pre-combusted (4 h at 450 °C) amber glass EPA vials (40 mL). Samples were acidified with 6 drops of concentrated HCl (12 mol L^−1^) to remove inorganic C, and stored in the dark at 4 °C until analysis. DOC concentrations were analyzed using a total organic C analyzer and applying the high-temperature catalytic oxidation method (TOC-VCSH; Shimadzu) modified from Sugimura and Suzuki (1988). Every 8–10 d the instrument was calibrated by measuring standard solutions of 0, 42, 83, 125, 208 and 417 *µ*mol C L^−1^, prepared from a potassium hydrogen phthalate standard (Merck 109017). Every measurement day, ultrapure (MilliQ) water was used to determine the instrument blank (< 1 *µ*mol C L^−1^). On every measurement day TOC analysis was validated with deep seawater reference (DSR) material provided by the Consensus Reference Materials Project of RSMAS (University of Miami) yielding values within the certified range of 42–45 *µ*mol C L^−1^. Additionally, two internal standards were prepared each measurement day using a potassium hydrogen phthalate (Merck 109017) with DOC concentration within the samples range. DOC of each sample was determined from 5–8 injections. The precision was < 4 % estimated as the standard deviation of replicate measurements divided by the mean.

Duplicate 1 mL samples for bacterioplankton and phytoplankton were fixed at a final concentration of 0.5 % glutaraldehyde for 15–30 min at 4 °C in the dark. After fixation, the samples were snap-frozen in liquid nitrogen and stored at -80 °C until further analysis.

Thawed samples were analyzed using a BD-FACSCalibur flow cytometer (*Becton Dickinson*, San Jose, Calif.) with a 15 mW air-cooled argon laser (Brussaard et al. 2004). Phytoplankton were enumerated for 10 min at 80 uL min^−1^ with the trigger on Chlorophyll a, red autofluorescence (Marie et al. 1999). Phycoerythrin containing cells (e.g. cyanobacterial *Synechococcus*) were discriminated by their orange autofluorescence. Bacterial samples were diluted in sterile TE-buffer, pH 8.0 (10 mmol L^−1^ Tris, *Roche Diagnostics*; 1 mmol L^−1^ EDTA, *Sigma-Aldrich*) to avoid electronic coincidence, and stained with nucleic acid-specific SYBR Green I to a final concentration of 1 x 10^−4^ of the commercial stock (Marie et al. 1999; Brussaard et al. 2004). Samples were corrected for blanks (TE-buffer with SYBR Green I) prepared and analyzed in a similar manner as the samples. Bacterial samples were incubated in the dark for 15 min at room temperature after which samples were allowed to cool down at room temperature. Samples were analyzed for 1 min at 40 *µ*L min^−1^. Listmode files were analyzed using CYTOWIN freeware (Vaulot et al. 1989).

### Sponge metrics

After the incubations, sponges were removed from their substrate and analyzed for volume (by water replacement) and (dripping) wet weight (WW). Then, all sponges were dried for 72 h in a drying oven at 60 °C to determine dry weight (DW). Randomly selected 1-cm^3^ cubes (*n* = 6) of each massive sponge were transferred into a pre-weighed crucible and combusted at 450 °C in a muffle furnace (4 h). Combusted samples were cooled to room temperature in a desiccator and weighed (ash weight). Subsequently, ash-free dry weight (AFDW) was calculated by subtracting ash weight from DW and normalized to total volume of the original sponge specimen. The rest of the dried sponges was crushed and ground up with mortar and pestle and stored in a desiccator until further analysis.

Samples for organic C content analysis were decalcified with 4 mol L^−1^ HCl to ensure removal of inorganic C and subsequently lyophilized for 24 h in a *FD5515 Ilchin Biobase* freeze-drier. After freeze-drying, aliquots of approximately 10 mg were placed in tin-capsules and analyzed on an Elemental Analyser (*Elementar Isotope cube, (Elementar GmbH*, Langenselbold, Germany) coupled to a BioVision isotope ratio mass spectrometer *(Elementar ltd*, Manchester, UK).

### Oxygen and carbon removal rates

To calculate changes in O_2_ concentrations over time, a linear regression analysis was performed for each individual incubation. Resulting net O_2_ removal rates were subsequently compared between sponge and seawater control incubations with a Welch’s *t*-test for each species and a respective set of seawater controls.

Initial net live bacterio- and phytoplankton removal rates were calculated assuming exponential clearance of cells in incubations (Scheffers et al. 2004; de Goeij et al. 2008). The live plankton fraction was dominated by two general cell types, heterotrophic bacteria and phytoplankton, the latter represented by *Synechococcus*-like cyanobacteria. To calculate net removal rates for each plankton component, the average initial cell concentrations of all incubations were used as a starting point. Standardized data were fitted to an inverse exponential model to calculate final cell concentrations. Final concentrations were subtracted from the initial corrected concentrations and differences were compared between treatments (sponge versus control incubations) using an unpaired t-test. Clearance rates (CR) were calculated according to Riisgård et al., 1993:

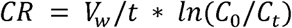

*V*_*w*_: water volume in incubation chamber (in mL)

*t*: duration of incubation (in min)

*C*_*0*_: initial cell (bacterial or phytoplankton) concentration (in *µ*mol C mL^−1^)

*C*_*t*_: cell concentration at time point t (in cells mL^−1^)

A conservative estimate of live particulate organic carbon (LPOC) removal was obtained using established conversion factors. Heterotrophic bacterial cells were converted using 30 fg C per bacterial cell (Fukuda et al. 1998; Leys et al. 2018) and phytoplankton using 470 fg C per *Synechococcus*-type cell (Bertilsson et al. 2003; Pile & Young 2006).

Initial net DOC removal rates were calculated by applying a 2G-model (de Goeij & van Duyl 2007; de Goeij et al. 2008). This is a simplified model to describe changes in DOC concentration over time, assuming that the complex and heterogeneous DOC pool comprises two major fractions: a fast-(C_f_) and slow-removable (C_s_) fraction, for labile and refractory components of DOM, respectively (de Goeij et al. 2008). In an assumed well-mixed system, the fast and slow removal fractions of DOC will be consumed according to their specific removal rate constants k_f_ and k_s_, respectively. The sum of the individual removal rates is used here to describe total DOC removal.

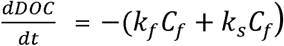

Integrating this equation yields the function that describes the concentration of DOC as a function of time:

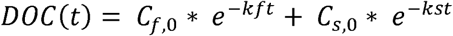

Experimental data is described with the model by estimating model variables C_f_ and C_s_ using a (10,000 iterations) minimalization routine (de Goeij et al. 2008). Initial DOC removal rate (the flux on t = 0) was calculated from the estimated values of these variables and is given by

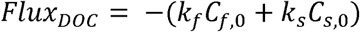

All fluxes of O_2_ and C in *µ*mol per different metric (sponge V, WW, DW, AFDW, or C content) were corrected for controls and incubation volume.

### Carbon mass balance

Total net organic carbon removal rates were estimated as the sum of net LPOC and DOC removal rates. O_2_ removal served as a proxy for respiration assuming a balanced molar ratio of carbon respiration to net O_2_ removal (1 mol C respired equals 1 mol O_2_ removed), yielding a respiratory quotient (RQ) of 1 (Hill et al. 2004; Yahel et al. 2003).

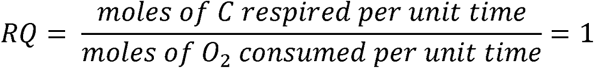

To establish a mass balance for the different deep-sea sponge species, the quotient ΔO_2_/ΔTOC was calculated using the exponential removal rates. Carbon budgets where only calculated for sponges of which we had complete sets of O_2_, LPOC and DOC data (*Vazella pourtalesi* (*n* = 4), *Geodia barretti* (*n* = 3), *Geodia atlantica* (*n* = 4), and *Acantheurypon spinispinosum* (*n* = 3).

## Results

### Sponge metrics

Sponge characteristics (phylogeny, growth form, abundance or associated microbes) and metric conversion factors are given in Table 1. Average sponge metrics (planar surface area, volume, WW, DW, AFDW, %C) are shown in Table S1. Encrusting sponges have a one-to two-orders of magnitude higher planar surface area to volume ratio (4.2–10.0) than massive sponges (0.2–0.3) and an order-of-magnitude higher volume to dry weight ratio (21.4–22.4 and 3.3–7.3, respectively). HMA sponges show a significantly higher organic C content than LMA sponges (*t* = -8.13, df = 27, *p* < 0.001), Table 1), with lowest values for the hexactinelid *V. pourtalesi*.

**Table 1.** Conversion factors between different standard metrics for six investigated deep-sea sponges. LMA = low microbial abundance sponges, HMA = high microbial abundance sponges. Planar surface area is the surface area covered in a 2D top view, volumes are measured by water displacement in mL and the weight is given as g dry weight (DW). Conversion factors are based on average sponge metrics (planar surface area, volume, wet weight (WW), DW, ash-free dry weight (AFDW), organic carbon (C) content) for all specimens used in the experiments shown in Table S1).

| Sponge species | Class | Growth Form | LMA/HMA | Planar surface area : Volume<br>(cm<sup>2</sup> : mL) | Volume : Weight<br>(mL : g dw) | organic C content<br>(% of dw) |
| --- | --- | --- | --- | --- | --- | --- |
| *Vazella pourtalesi* | Hexactinellid | Massive, vase | LMA | 0.3 | 5.2 | 5.5 |
| *Geodia barretti* | Demosponge | Massive, ball | HMA | 0.3 | 3.3 | 15.9 |
| *Geodia atlantica* | Demosponge | Massive | HMA | 0.3 | 7.3 | 20.3 |
| *Craniella zetlandica* | Demosponge | Massive, ball | HMA | 0.2 | 4.3 | 20.4 |
| *Hymedesmia paupertas* | Demosponge | Encrusting, sheet | LMA | 10.0 | 21.4 | 12.6 |
| *Acantheurypon spinispinosum* | Demosponge | Encrusting, sheet | LMA | 4.2 | 22.4 | 10.9 |

### Oxygen removal rates

The concentration of O_2_ in the incubation chambers linearly decreased with time for *V. pourtalesi* (*t* = 4.59, df = 7, *p* < 0.01), *G. barretti* (*t* = 3.69, df = 11, *p* < 0.01), *G. atlantica* (*t* = 5.11, df = 5, *p* < 0.01), *C. zetlandica* (*t* = 3.5, df = 3, *p* < 0.05), *H. paupertas* (*t* = 4.38, df = 2, *p* < 0.05) and *A. spinispinosum*, (*t* = 7.96, df = 5, *p* < 0.001) compared to seawater control incubations. Average O_2_ removal rates per species are depicted in Table 2. Examples of O_2_ concentration profiles during incubations for all species and controls are shown in figure S1. O_2_ removal rates for all sponges averaged 3.3. ± 2.8 *µ*mol O_2_ DW_sponge_ h^−1^ (mean ± SD throughout text unless stated otherwise), ranging from 1.0 (*C. zetlandica*) to 7.8 (*A. spinispinosum*).

**Table 2.**
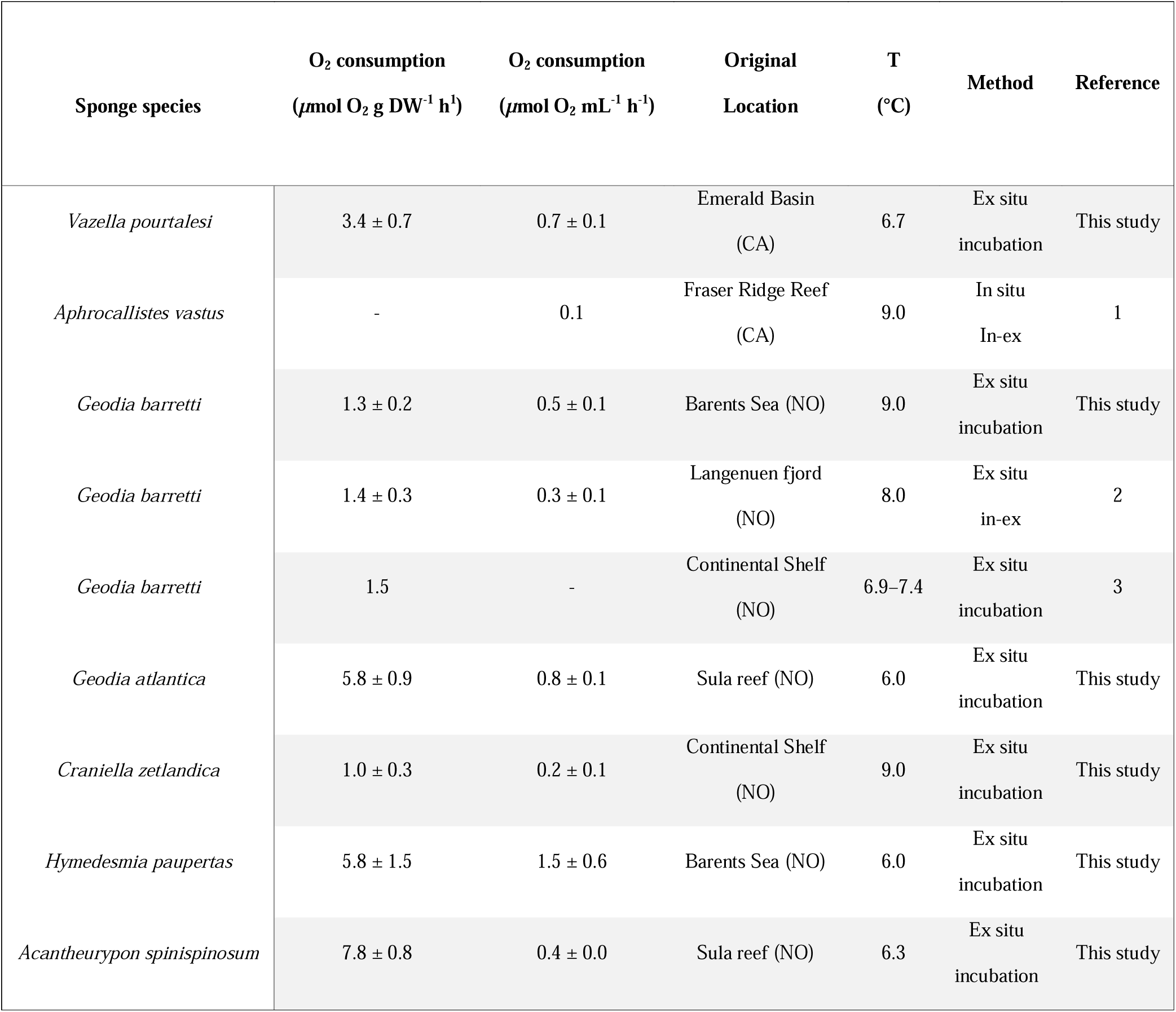
Oxygen consumption by deep-sea sponge species (mean ± SE). CA = Canada, NO = Norway. (1) Leys et al., 2011, (2) Leys et al. 2018, (3) Kutti et al. 2013.

### LPOC removal rates

Bacterioplankton concentrations decreased in incubations with *G. barretti* (*t* = 2.44, df = 19, *p* < 0.05), *V. pourtalesi* (*t* = 5.91, df = 9, *p* < 0.001), *G. atlantica* (*t* = 6.62, df = 5, *p* < 0.01), *H. paupertas* (*t* = 2.81, df = 4, *p* < 0.05) and *C. zetlandica* (*t* = 4.25, df = 8, *p* < 0.01) compared to seawater control incubations (Fig. 3A–E). Incubations with *A. spinispinosum* showed no significant decrease in bacterioplankton compared to control incubations (*t* = -0.72, df = 4, *p* = 0.51) (Fig. 3F). Average bacterial C removal and clearance rates (CR) per species are presented in Table 3. Bacterial C removal rates averaged 0.25 ± 0.35 *µ*mol C DW_sponge_ h^−1^ for all species, ranging between 0.00 (*A. spinispinosum*) and 0.82 (*V. pourtalesi*) (Table 3). Bacterial CRs averaged 0.69 ± 1.06 mL mL^−1^ min-^1^ for all species, ranging from 0.00 (*A. spinispinosum*) to 2.22 (*V. pourtalesi*).

**Table 3.**
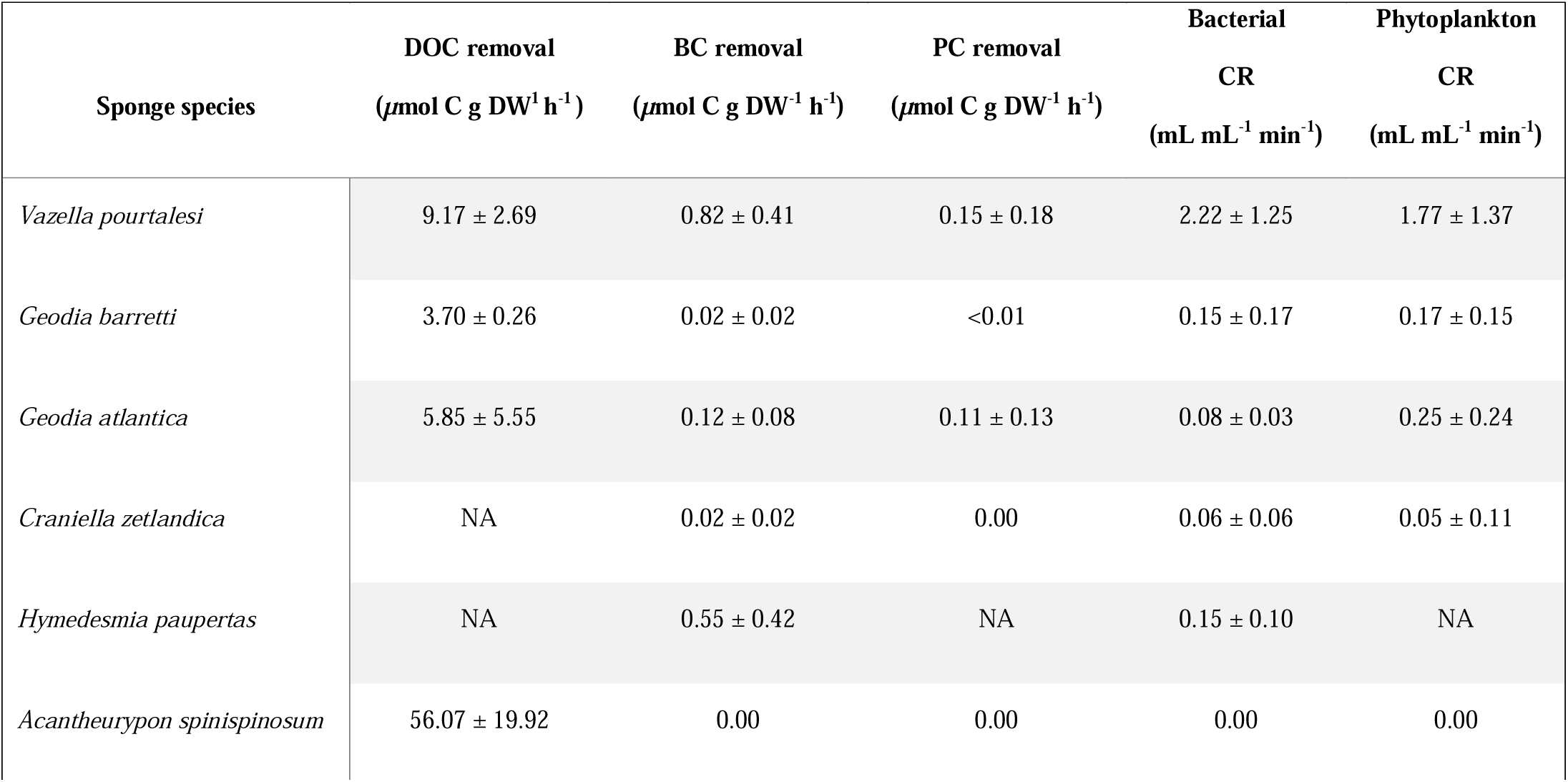
Average (± SD) net dissolved organic carbon (DOC), bacterio- and phytoplankton carbon (BC and PC, respectively) removal rates, and bacterio- and phytoplankton clearance rates per sponge species. Net removal rates for bacterio- and phytoplankton are based on exponential uptake during incubations, whereas net removal rates for DOC are based on a 2G-model uptake. DOC = dissolved organic carbon, BC = bacterial carbon, PC = phytoplankton, CR = clearance rate.

| Sponge species | DOC removal<br>($\mu\text{mol C g DW}^{-1} \text{h}^{-1}$) | BC removal<br>($\mu\text{mol C g DW}^{-1} \text{h}^{-1}$) | PC removal<br>($\mu\text{mol C g DW}^{-1} \text{h}^{-1}$) | Bacterial<br>CR<br>($\text{mL mL}^{-1} \text{min}^{-1}$) | Phytoplankton<br>CR<br>($\text{mL mL}^{-1} \text{min}^{-1}$) |
| --- | --- | --- | --- | --- | --- |
| *Vazella pourtalesi* | 9.17 ± 2.69 | 0.82 ± 0.41 | 0.15 ± 0.18 | 2.22 ± 1.25 | 1.77 ± 1.37 |
| *Geodia barretti* | 3.70 ± 0.26 | 0.02 ± 0.02 | <0.01 | 0.15 ± 0.17 | 0.17 ± 0.15 |
| *Geodia atlantica* | 5.85 ± 5.55 | 0.12 ± 0.08 | 0.11 ± 0.13 | 0.08 ± 0.03 | 0.25 ± 0.24 |
| *Craniella zetlandica* | NA | 0.02 ± 0.02 | 0.00 | 0.06 ± 0.06 | 0.05 ± 0.11 |
| *Hymedesmia paupertas* | NA | 0.55 ± 0.42 | NA | 0.15 ± 0.10 | NA |
| *Acantheurypon spinispinosum* | 56.07 ± 19.92 | 0.00 | 0.00 | 0.00 | 0.00 |

**Figure 3.**
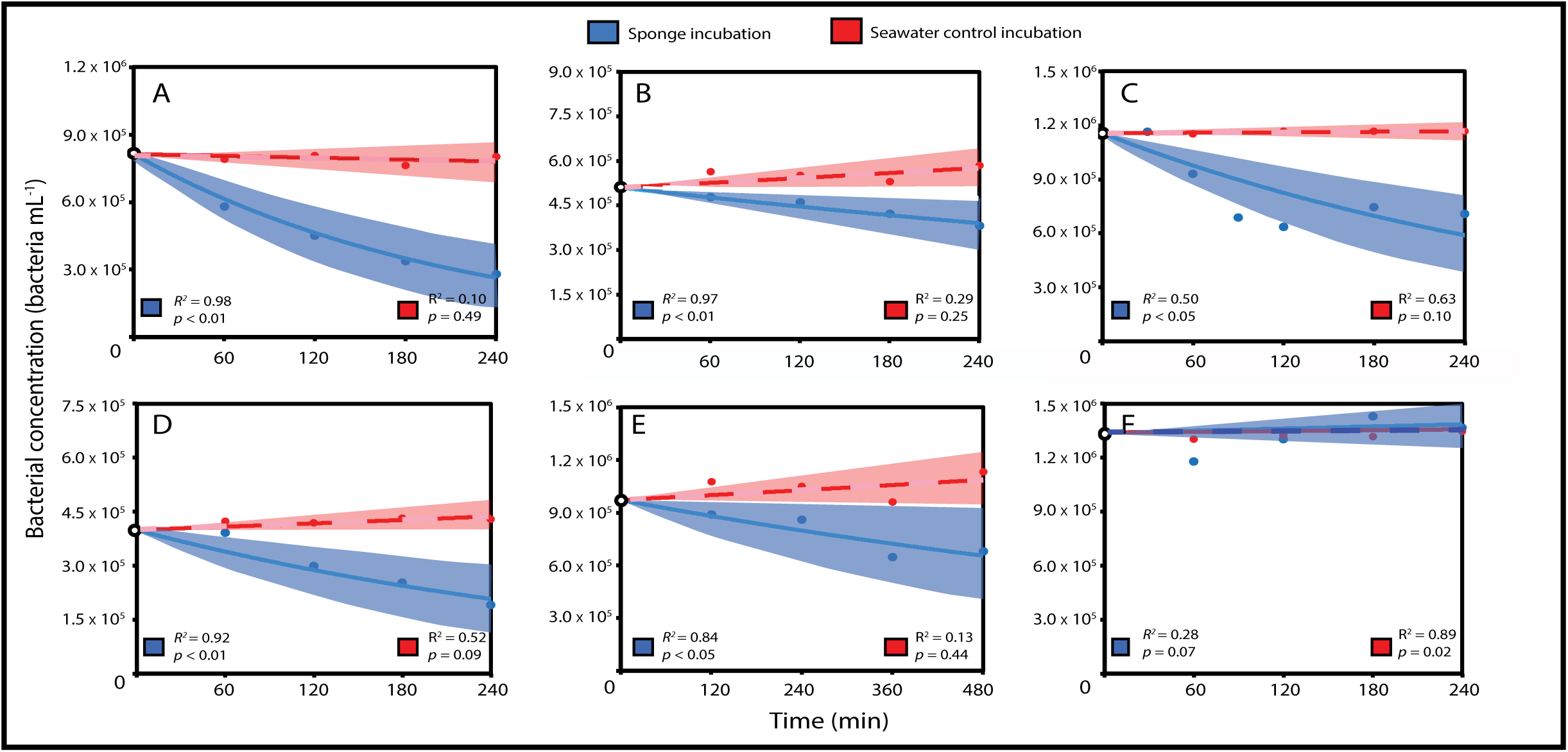
Average abundances of bacterioplankton during incubations with six deep-sea sponge species in comparison to seawater controls incubations over time. Sponges (blue) versus seawater controls (red). A: *Vazella pourtalesi* (*n* = 7), B; *Geodia barretti* (*n* = 12), C; *Geodia atlantica* (*n* = 6), F: D: *Craniella zetlandica* (*n* = 4), E: *Hymedesmia paupertas* (*n* = 3), F: *Acantheurypon spinospinosum* (*n* = 4). Bacterial decrease is modelled with an exponential fit, shades depict 95% confidence intervals of the model. Note that x- and y-axis show different ranges per species.

Compared to control incubations, phytoplankton (i.e. S*ynechococcus*-type cyanobacteria) concentrations decreased in incubations with *V. pourtalesi* (*t* = 5.34, df = 9, *p* < 0.001), *G. barretti* (*t* = 2.20, df = 11, *p* < 0.05) and *G. atlantica* (*t* = 11.92, df = 6, *p* < 0.001). Incubations with *C. zetlandica* (*t* = 1.23, df = 3, *p* = 0.31) and *A. spinispinosum* (*t* = 1.56, df = 7, *p* = 0.16) showed no significant decrease compared to seawater control incubations. Average phytoplankton C fluxes per species are presented in Table 3.

Phytoplankton C removal rates averaged 0.04 ± 0.07 *µ*mol C DW_sponge_ h^−1^ for all species, ranging from 0.00 (*A. spinispinosum/C. zetlandica*) to 0.15 (*V. pourtalesi*) (Table 3). Phytoplankton CRs averaged 0.54 ± 0.96 mL mL^−1^ min-^1^ for all species, ranging between 0.00 (*A. spinispinosum*) and 1.77 (*V. pourtalesi*).

Combined plankton removal rates amounted to total LPOC uptake rates of, on average, 0.30 ± 0.39 *µ*mol C DW_sponge_ h^−1^, ranging from 0.00 (*A*. spinispinosum) to 0.97 (*V. pourtalesi*).

### DOC removal rates

DOC concentration for incubations with four different species: *V. pourtalesi* (*n* = 4), *G. barretti* (*n* = 3), *G. atlantica* (*n* = 4), and *A. spinispinosum* (*n* = 3) fitted the 2G model and thereby showed significant removal of DOC (Fig. 4 and Supplementary Fig. S2).

**Figure 4.**
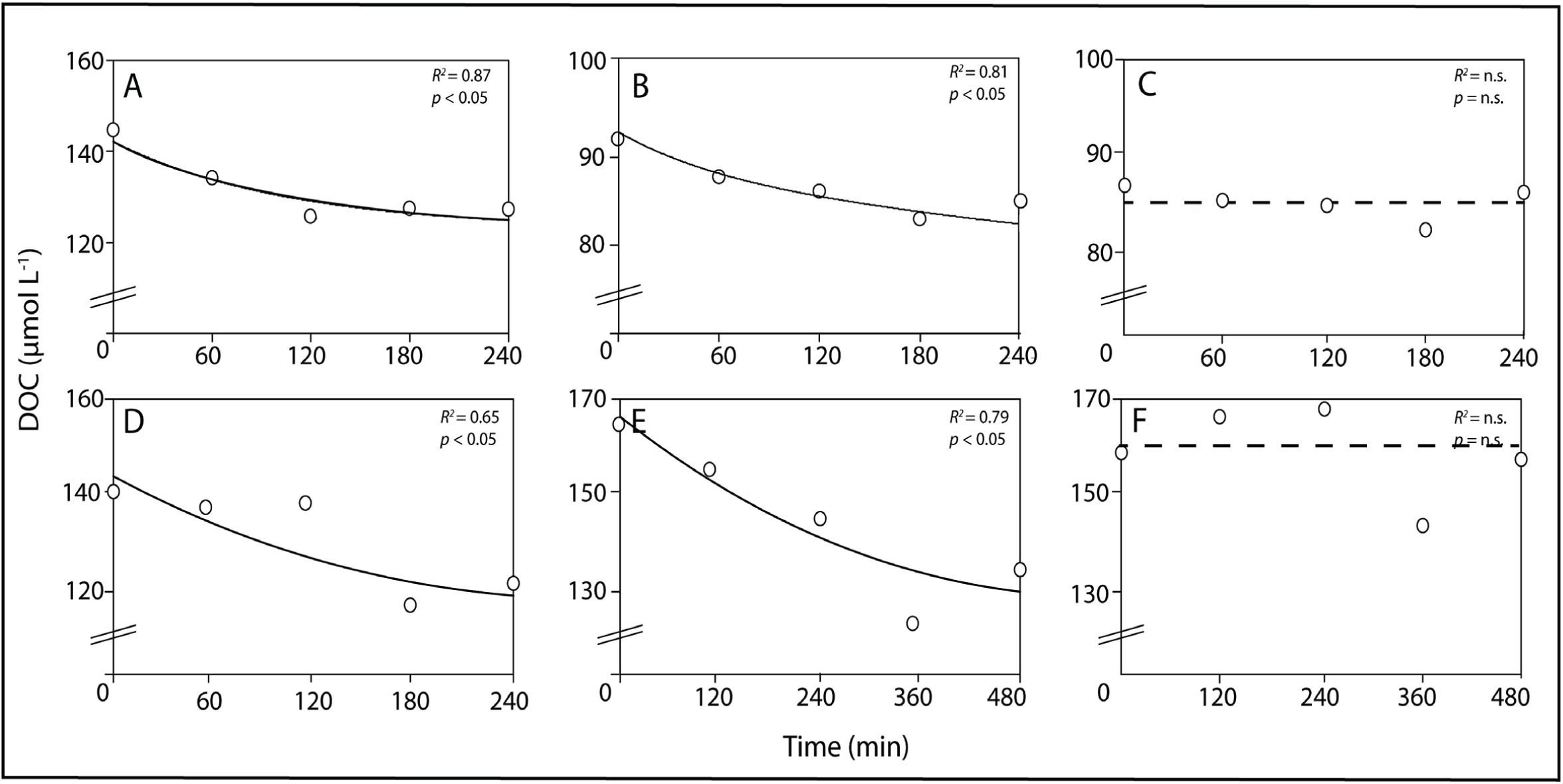
DOC removal over time by four deep-sea sponge sponge species compared to seawater controls in *ex situ* incubations. A; *Vazella pourtalesi* B: *Geodia barretti* C: Seawater control, D: *Geodia atlantica*, E: *Acantheurypon spinispinosum*, F: Seawater control. Trend lines are given by a 2G model fit.

Unfortunately, some time-series could not be analyzed due to technical difficulties. No DOC removal occurred in the seawater controls. DOC removal rates averaged 18.70 ± 25.02 *µ*mol C DW_sponge_ h^−1^ for all sponges, ranging from 3.70 (*G*. barretti) to 56.07 (*A. spinispinosum*) (Table 3).

### Carbon mass balance

Mass balances, constructed for the four species where both LPOC and DOC were measured, showed that more than 90% of the average net TOC removal was accounted for by DOC (*V. pourtalesi* 92.0 ± 5.5 %, *G. barretti* 99.5 ± 0.5 %, *G. atlantica* 93.6 ± 8.4 %, *A. spinospinosum* 100 %) (Table 4). All species except *A. spinospinosum* also removed LPOC from the water, yet this organic carbon source accounted for less than 10 % of the net total TOC removal. Assuming a RQ of 1 in combination with exponential removal of LPOD and DOC during the incubations, we find all four species can match their minimal required carbon uptake (ΔO_2_/ΔTOC of 1.0 or lower), but only when DOC is included in the mass balance. Both HMA species show higher ΔO_2_/ΔTOC values than the two LMA species.

**Table 4.** Carbon mass balance for four deep-sea sponge species. The mass balance was based on linear fluxes of oxygen uptake, exponential net removal rates of bacterio- and phytoplankton organic carbon (LPOC) and 2G-model exponential net removal rates for dissolved organic carbon (DOC). Net total organic carbon (TOC) removal rates are calculated as the sum of LPOC and DOC.

| Sponge species | O<sub>2</sub><br>($\mu\text{mol h}^{-1} \text{ g DW}^{-1}$) | TOC<br>($\mu\text{mol h}^{-1} \text{ g DW}^{-1}$) | Exponential mass balance<br>$\Delta\text{O}_2/\Delta\text{TOC}$ |
| --- | --- | --- | --- |
| *Vazella pourtalesi* | $3.19 \pm 1.96$ | $9.87 \pm 2.36$ | 0.32 |
| *Geodia barretti* | $1.93 \pm 1.09$ | $2.97 \pm 1.87$ | 0.65 |
| *Geodia atlantica* | $4.76 \pm 1.76$ | $6.07 \pm 5.55$ | 0.78 |
| *Acantheurypon spinispinosum* | $8.44 \pm 1.04$ | $56.07 \pm 19.92$ | 0.15 |

## Discussion

This is the first study that combines measured uptake rates of dissolved oxygen (O_2_), dissolved organic carbon (DOC) and live planktonic particulate organic carbon (LPOC) by multiple deep-sea sponges. We found that for the four investigated species where both LPOC and DOC were measured, DOC accounted for 92–100 % of the total organic carbon (TOC) uptake. Only when DOC is included as organic carbon source, these deep-sea sponges surpass their minimal respiratory carbon demands. Furthermore, metabolic rates, morphometrics, and conversion factors for six dominant North-Atlantic deep-sea sponge species are presented.

### Oxygen and carbon fluxes

O_2_ removal rates per DW of deep-sea sponges show consistency throughout literature, as most are roughly within the same order of magnitude (equal or less than factor 10 difference; Table 2). For *G. barretti*, O_2_ removal rates found in this study are closely comparable with other rates reported, regardless of the method used (Kutti et al. 2013; Leys et al. 2018). When comparing respiration rates of deep-sea sponges to those reported for temperate (e.g., Thomassen & Riisgard 1995; Coma et al. 2002) and tropical sponges (e.g., Reiswig 1974; Yahel et al. 2003), rates of deep-sea sponges are consistently one to two orders-of-magnitude lower. Correspondingly, POC and DOC removal rates of deep-sea sponges are lower than those found for tropical and temperate species (e.g., de Goeij et al. 2008; Hoer et al. 2018).

As depicted in figure 5, differences in O_2_ and POC removal rates can be explained by the positive effect of temperature on metabolism and physiological processes (see also Clarke and Fraser 2004). DOC removal rates seem to follow a similar trend, yet due to the very limited amount of data available, the relation with temperature was not found to be significant. Remarkably, O_2_, POC and (specifically) DOC fluxes appear to be higher for encrusting sponges compared to massive growth forms (Fig. 5). For example, the deep-sea encrusting sponge *A. spinispinosum* has an order-of-magnitude higher DOC flux than massive deep-sea species (56.1 *µ*mol C h^−1^ g DW^−1^ versus 3.7–9.2 *µ*mol C h^−1^ g DW^−1^), as is the case for encrusting tropical species (218.3–253.3 *µ*mol C h^−1^ g DW^−1^, de Goeij et al. 2008) compared to massive tropical species (10.0–11.9 *µ*mol C h^−1^ g DW^−1^, Yahel et al. 2003, Hoer et al. 2018). This corroborates earlier suggestions that high surface-to-volume ratios enable encrusting sponges to have higher retention and removal efficiencies compared to massive species (Abelson 1993; Kötter 2003; de Goeij et al. 2017).

**Figure 5.**
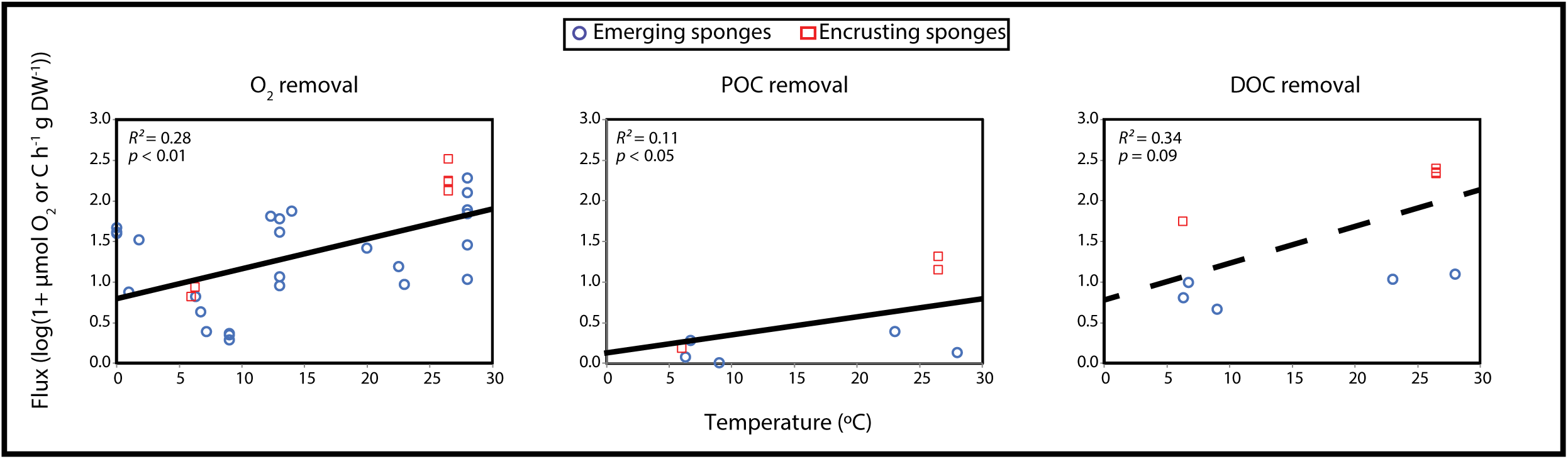
Oxygen and particulate - and dissolved carbon fluxes for tropical, temperate, and cold-water marine sponges plotted against temperature. Fluxes are log transformed. Red squares depict encrusting sponges, blue circles depict emerging/massive sponges. R^2^ values are based on the linear regression of all values (encrusting + emerging). Regression lines are given by log(1+ *µ*mol O2 h^−1^ g DW^−1^) = 0.033 * T(°C) + 0.90, (log(1+ *µ*mol POC h^−1^ g DW^−1^) = 0.017 * T(°C) + 0.21 and (log(1+ *µ*mol DOC h^−1^ g DW^−1^) = 0.041 * T(°C) + 0.77. All non-transformed fluxes are given in table S2.

In addition to morphology, higher net DOC removal rates are generally predicted for HMA sponges in comparison with LMA sponges, as microbes are considered to play an important role in the processing of DOM (Reiswig 1974; Freeman & Thacker 2011; Maldonado et al. 2012; Hoer et al. 2018). However, both LMA species used in this study, *A. spinispinosum* and *V. pourtalesi*, showed high uptake rates of DOC (56.1 and 9.2 *µ*mol C h^−1^ g DW^−1^, respectively), despite their different growth forms (encrusting versus massive) and different phylogeny (demosponge versus hexactinellid). Interestingly, other hexactinellids were previously not found to consume DOM (Yahel et al. 2007), where in this study *V. pourtalesi* shows the second highest DOC removal rates and DOC accounts for 90 % of its TOC uptake. These difference could imply species-too-species differences or geographical differences in organic carbon availability. However, Yahel et al. (2007) did not directly measure DOC, but derived it from TOC analysis, potentially resulting in an underestimation of actual DOC removal rates. Our results thereby add to the increasing body of evidence that also sponges with low microbial abundances are capable of consuming DOC (de Goeij et al. 2013; Rix et al. 2016, 2017; Morganti et al. 2017).

### Interpretation of sponge metabolic rates

Our understanding and interpretation of metabolic rates at both organism and ecosystem scale is currently hampered by two issues. Firstly, a general lack of available oxygen and (predominantly dissolved organic) carbon fluxes of deep-sea species, but also for marine sponges in general (discussed in de Goeij et al. 2017). Specifically, no data are available on encrusting HMA species, which raises the question whether the high fluxes of encrusting LMA sponges are due to their encrusting morphology or due to their limited abundances of symbionts. Secondly, the use of a multitude of metrics (e.g., m^−2^, WW, DW, AFDW) in combination with a lack of conversion factors, makes it almost impossible to compare metabolic rates between different sponge species. Yet, this is crucial to upscale fluxes from organism to ecosystem level in order to elucidate the role of marine sponges in biogeochemical cycles and overall ecosystem functioning. But, which metric should be used? In general, this may depend on the context and the research question at hand. For example, when extrapolating individual fluxes to the ecosystem level, planar surface area is potentially the most practical standardization metric in use (read: fast and low-cost). Abundance data in deep-sea —but also in shallow-water— habitats are usually collected via 2D video surveys or photo quadrants using ROV’s (van Soest 2007; Roberts et al. 2009). However, 2D planar surface area severely underestimates the volume and (organic) biomass of erect versus flat organisms (e.g., massive versus encrusting sponges; see also discussion in de Goeij et al. 2017). Wet weight is another commonly used metric, but the most subjective, since the loss of weight in time before measuring can significantly change within and between species. Arguably the best metric to standardize metabolic rate is organic biomass (i.e. AFDW) or organic carbon content, excluding ecologically inert hard constituents, such as silica spicules (Rutzler, 1978). However, an increase in inorganic spicule content requires additional energetic costs at the expense of organic material (McDonald et al. 2002; Carballo et al. 2006). Therefore, in ecological terms, volume and DW provide alternatives. Volume, similar to WW, compromised by effects of large variations in shape, form and tissue densities and compositions of sponges (Diaz & Rutzler, 2001). We therefore use DW here as comparative measure and suggest that future physiological studies on sponges best provide a combination of metrics on volume, area cover, DW, AFDW, elemental composition (e.g., C or N content) and/or conversion factors between these metrics.

### Deep-sea sponge carbon budgets

The contribution of DOC to the TOC uptake of the investigated sponges (92–100 %) is at the high end of the range reported for shallow water sponges (56–97; see Table 1 in de Goeij et al. 2017). Indirect measurements recently suggested that DOC accounts for 95 % of the TOC uptake of *G. barretti* (Leys et al. 2018), which is very close to the fluxes presented here. Although mass balances for all species where LPOC and DOC were measured are largely positive (table 4), both HMA species show higher ΔO_2_/ΔTOC values than the two LMA species. These differences might be explained by aerobic microbial processes in HMA sponges, such as nitrification (Hoffmann et al. 2009) or ammonia oxidation (Mohamed et al. 2010), which require O_2_ in addition to the O_2_ demand based on carbon respiration. Moreover, the organic carbon uptake needed to balance respiration requirements of HMA sponges is potentially further reduced by sponge-associated chemoautotrophs using inorganic carbon sources which are transferred to the sponge host (van Duyl et al. 2008; Pita et al. 2018; Shih et al. 2019). A second issue with the calculating mass balances is the use of RQ value. The value of 1 used in this study applies to the oxidation of simple sugars (CH_2_O)_x_. In reality, proteins and nucleic acids have RQ values ranging from 0.71–0.83 (Kleiber, 1975), meaning that depending on the macromolecules respired, less than 1 mole C could be needed to balance consumption of 1 mole O, further reducing the amount of C needed to balance respiration requirements. In addition to respiration, other processes such as growth, cell-turnover and release, reproduction, and the production of metabolites require organic carbon. Therefore, a complete carbon budget should include these processes. However, deep-sea sponges most likely grow slowly (Leys & Lauzon 1998), and we assume that within the short (2–8 h) timeframe of our incubations, growth is negligible. For several shallow water encrusting sponges, a rapid cell turnover and the subsequent release of “old” cells as detritus was shown (de Goeij et al. 2009, 2013; Alexander et al. 2014; Rix et al. 2016). This “loss of carbon” could have a major impact on carbon budgets. However, Leys et al. (2018) reported no production of new cells during experiments with *G. baretti*, suggesting minimal investment in cell turnover in the investigated time frame. In contrast, Rix and colleagues (2016) found that the deep-sea encrusting sponge *Hymedesmia coriacea* converted 39 % of organic carbon derived from deep-sea coral mucus into detritus, and detritus production by deep-sea sponges has been argued to have a major contribution to the total sedimentation rate of the Greenland–Iceland–Norwegian seas (Witte et al. 1997). Concludingly, reports on deep-sea sponge detritus production and cell turnover are contradictive and still very limited, which does not warrant generalizations at this point. Likewise, only limited data is available on the reproduction of deep-sea sponges (Spetland et al. 2007) as well as seasonal changes in metabolic rates (Morley et al. 2016). Lastly, particularly HMA sponges, such as *G. barretti*, are known to produce secondary metabolites for chemical defense against surface settlers and grazers (Hedner et al. 2006; Sjo□gren et al. 2011). However, to our knowledge no studies have been performed on the metabolic costs of this metabolite production.

## Conclusion

In this study we showed for the first time that multiple deep-sea sponge species are capable of consuming natural DOC, and that this DOC is essential to satisfy their minimal respiratory carbon demand. However, although bacterio- and phytoplankton contributed only a small fraction (< 10 %) to the TOC uptake, these particulate food sources may contain valuable nutrients, such as vitamins, fatty acids, and amino acids (Putter 1925; Phillips 1984), which are essential for anabolic processes. We therefore hypothesize that DOC comprises the main carbon source for deep-sea sponges to sustain their minimal energetic requirements. But the supplementation with bacterio- and phytoplankton and possibly detritus, particularly during episodic food pulses after phytoplankton blooms, is essential to support anabolic processes such as somatic growth, reproduction, and cell turnover. The effective consumption of both dissolved and particulate food therefore allows deep-sea sponges to thrive in otherwise food-limited environments.

## Acknowledgements

We thank all our collaborators at the EU Horizon 2020 SponGES project; Dr. Ellen Kenchington at the Bedford Institute of Oceanography (BIO), Nova Scotia, Canada, and the Department of Biological Sciences at the University of Bergen, Norway, for the use of facilities and equipment. Many thanks to the ROV crews of both the ÆGIR 6000 in Norway and the Deep Sea Systems International Oceaneering in Canadian waters, for their careful collection of the sponges. DFO provided the ship time on the CCGS *Hudson* for the collection of the *Vazella pourtalesi* samples and we thank Dr. Ellen Kenchington for collecting the samples at sea under less than ideal conditions. We further thank Dr. L. Woodall, NEKTON, for providing the ROV for the collection of the *V. pourtalesi*. Thanks to Erik Wurz for his help maintaining the onboard aquaria. We also thank Jorien Schoorl and Rutger van Hall for their technical assistance at the University of Amsterdam, Niklas Kornder and Dr. Emiel van Loon for their advice on the statistical analysis, Heikki Savolainen for his technical support at the University of Bergen, Dr. Alice Ortmann (BIO), Anna Noordeloos (NIOZ) and Santiago Gonzalez (NIOZ) for their help with the flow cytometry analysis, and Jon Roa at GEOMAR for the analysis of the DOC samples. This project has received funding from the European Research Council under the European Union’s Horizon 2020 research and innovation programme (SponGES grant agreement n° 679849 and ERC starting grant agreement n° 715513 to J.M. de Goeij). Additionally, this project has received funding from the KNAW Academy Ecology fund in 2017 and 2018 (personal grant to M.C. Bart). Salary for Gabrielle Tompkins was funded through the Canadian Department of Fisheries and Oceans (DFO) International Governance Strategy internal funding awarded to Dr. Ellen Kenchington (DFO, BIO). This document reflects only the authors’ views and the Executive Agency for Small and Medium-sized Enterprises (EASME) is not responsible for any use that may be made of the information it contains.

