## Supplementary Table S1 for "Dissolved organic carbon (DOC) is essential to balance the metabolic demands of North-Atlantic deep-sea sponges"

**Table S1. Average morphometrics for all species used in the incubation experiments** (mean ± SD).

| **Species** | **n** | **Planar surface area (cm^2^)** | **Volume (mL)** | **WW (g)** | **DW (g)** | **AFDW (g)** | **Organic C content (%)** |
| --- | --- | --- | --- | --- | --- | --- | --- |
| Vazella pourtalesi | 7 | 4.4 ± 3.3 | 14.7 ± 12.4 | 28.2 ± 24.9 | 2.8 ± 2.3 | 0.5 ± 0.4 | 5.5 ± 0.8 |
| Geodia barretti | 12 | 20.6 ± 13.8 | 79.7 ± 69.2 | 99.7 + 71.3 | 24.4 ± 18.2 | 10.8 ± 8.0 | 15.9 ± 2.4 |
| Geodia atlantica | 6 | 86.0 ± 38.1 | 308.3 ± 106.8 | 438.3 ± 139.3 | 42.2 ± 15.8 | 21.5 ± 8.1 | 20.3 ± 2.8 |
| Craniella zetlandica | 4 | 41.7 ± 11.2 | 232.5 ± 73.3 | 242.9 ± 72.3 | 54.7 ± 17.3 | 24.6 ± 7.8 | 20.3 ± 3.3 |
| Hymedesmia paupertas | 3 | 42.0 ± 6.1 | 4.2 ± 0.6 | 1.0 ± 0.7 | 0.2 ± 0.2 | - | 12.6 ± 1.8 |
| Acantheurypon spinospinosum | 4 | 165.8 ± 26.7 | 45.0 ± 9.7 | 8.2 ± 1.0 | 2.0 ± 0.3 | 0.7 ± 0.1 | 10.9 ± 0.8 |
