## Supplementary Table S2 for "Dissolved organic carbon (DOC) is essential to balance the metabolic demands of North-Atlantic deep-sea sponges"

**Table S2. Oxygen (O_2_), Particulate organic carbon (POC) and dissolved organic carbon (DOC) fluxes in µmol DW^-1^ h^-1^ of various sponge species from literature**. HMA = high microbial, LMA = low microbial, T = temperature (°C), O_2_, POC and DOC fluxes in µmol h^-1^ g dw_sponge_^-1^. 1 = Witte & Graf 1996, 2 = Kowalke 2000, 3 = Kutti et al. 2013, 4 =Leys et al. 2018, 5 = Cotter 1978, 6 = Murray 2009, 7 = Tomassen & Riisgard 1995, 8 = Coma 2002, 9 = Hadas et al. 2008, 10 = Yahel et al. 2003, 11 = de Goeij et al. 2013 unpublished data, 12 = de Goeij et al. 2008, 13 = Reiswig 1974, 14 = Hoer et al. 2018.

| **Sponge** | **Class** | **Growth form** | **HMA/LMA** | **T** | **O_2_** | **POC** | **DOC** | **Ref.** |
| --- | --- | --- | --- | --- | --- | --- | --- | --- |
| Craniella cranium | Demosponge | Emerging | HMA | 0 | 40.0 | n.a. | n.a. | 1^ᵻ^ |
| Thenea abyssorum | Demosponge | Emerging | n.a. | 0 | 47.0 | n.a. | n.a. | 1 |
| Thenea muricate | Demosponge | Emerging | n.a. | 0 | 41.5 | n.a. | n.a. | 1 |
| Isodictia kerguelensis | Demosponge | Emerging | n.a. | 1.0 | 6.7 | n.a. | n.a. | 2^#^ |
| Mycale acerata | Demosponge | Emerging | LMA | 1.8 | 32.8 | n.a. | n.a. | 2*^,##^ |
| Hymedesmia paupertas | Demosponge | Encrusting | LMA | 6.0 | 5.8 | 0.55 | n.a. | This study |
| Geodia atlantica | Demosponge | Emerging | HMA | 6.3 | 5.8 | 0.21 | 5.56 | This study |
| Acantheurypon spinospinosum | Demosponge | Encrusting | LMA | 6.3 | 7.8 | n.a. | 56.07 | This study |
| Vazella pourtalesi | Hexactinellid | Emerging | LMA | 6.7 | 3.4 | 0.97 | 9.2 | This study |
| Geodia barretti | Demosponge | Emerging | HMA | 7.2 | 1.5 | n.a. | n.a. | 3 |
| Geodia barretti | Demosponge | Emerging | HMA | 8.0 | 1.4 | 0.04 | n.a. | 4 |
| Geodia barretti | Demosponge | Emerging | HMA | 9.0 | 1.3 | 0.02 | 3.70 | This study |
| Craniella zetlandica | Demosponge | Emerging | HMA | 9.0 | 1.0 | 0.02 | n.a. | This study |
| Sycon ciliatum | Calcareous | Emerging | LMA | 13.0 | 65.8 | n.a. | n.a. | 5^+^ |
| Stellata sp. | Demosponge | Emerging | HMA | 13.0 | 10.9 | n.a. | n.a. | 6^+^ |
| Tethya bergquistae | Demosponge | Emerging | LMA | 13.0 | 8.3 | n.a. | n.a. | 6* |
| Mycale sp. | Demosponge | Emerging | LMA | 13.0 | 61.1 | n.a. | n.a. | 6* |
| Leucosolenia echinata | Calcareous | Emerging | LMA | 14.0 | 40.9 | n.a. | n.a. | 6^+^ |
| Halichondia panicea | Demosponge | Emerging | LMA | 20.0 | 75.1 | n.a. | n.a. | 7* |
| Dysidea avarata | Demosponge | Emerging | LMA | 22.5 | 25.8 | n.a. | n.a. | 8* |
| Negombata magnifica | Demosponge | Emerging | n.a. | 23.0 | 14.9 | n.a. | n.a. | 9 |
| Theonella swinhoei | Demosponge | Emerging | HMA | 26.5 | 8.6 | 1.5 | 1.6 | 10 ^‡^ |
| Chondrilla caribensis | Demosponge | Encrusting | HMA | 26.5 | 181.0 | n.a. | n.a. | 11 |
| Halisarca caerulea | Demosponge | Encrusting | LMA | 26.5 | 336.0 | 20.0 | 218.3 | 11 |
| Mycale microsigmatosa | Demosponge | Encrusting | LMA | 26.5 | n.a. | 20.0 | 253.3 | 11 |
| Merlia normani | Demosponge | Encrusting | LMA | 26.5 | n.a. | 13.3 | 226.7 | 12 |
| Scopalina ruetzleri | Demosponge | Encrusting | LMA | 26.5 | 134.0 | n.a. | n.a. | 11 |
| Haliclona implexiformis | Demosponge | Encrusting | LMA | 26.5 | 173.7 | n.a. | n.a. | 11 |
| Verongia gigantea | Demosponge | Emerging | HMA | 28.0 | 127.5 | n.a. | n.a. | 13^###,‡‡^ |
| Hyrtios sp. | Demosponge | Emerging | HMA | 28.0 | 78.7 | n.a. | n.a. | 6* |
| Xestospongia muta | Demosponge | Emerging | HMA | 28.0 | 10.0 | 0.4 | 1.9 | 14^•^ |
| Tethya crypta | Demosponge | Emerging | LMA | 28.0 | 28.6 | n.a. | n.a. | 13^###,‡‡^ |
| Dysidea herbacea | Demosponge | Emerging | LMA | 28.0 | 69.8 | n.a. | n.a. | 6* |
| Mycale sp. | Demosponge | Emerging | LMA | 28.0 | 196.6 | n.a. | n.a. | 13^###,‡‡,^ * |

ᵻ Also named *Tethya cranium*, # Ash content based on Morley (2016), ## Ash content based on McClinktock (1987), * HMA/LMA based on Moutinho-Silva (2017), + HMA/LMA based on Vacelet & Donaday (1977), ‡Temperature from on Yahel (2002), ### Ash content from Reiswig (1971a), ‡‡ Temperature from Osinga et al. (1999), • Conversion volume – dry weight from Fiore et al. (2013).
