## Supplementary figures and images for "Dissolved organic carbon (DOC) is essential to balance the metabolic demands of North-Atlantic deep-sea sponges"

### Supplementary Figure S1

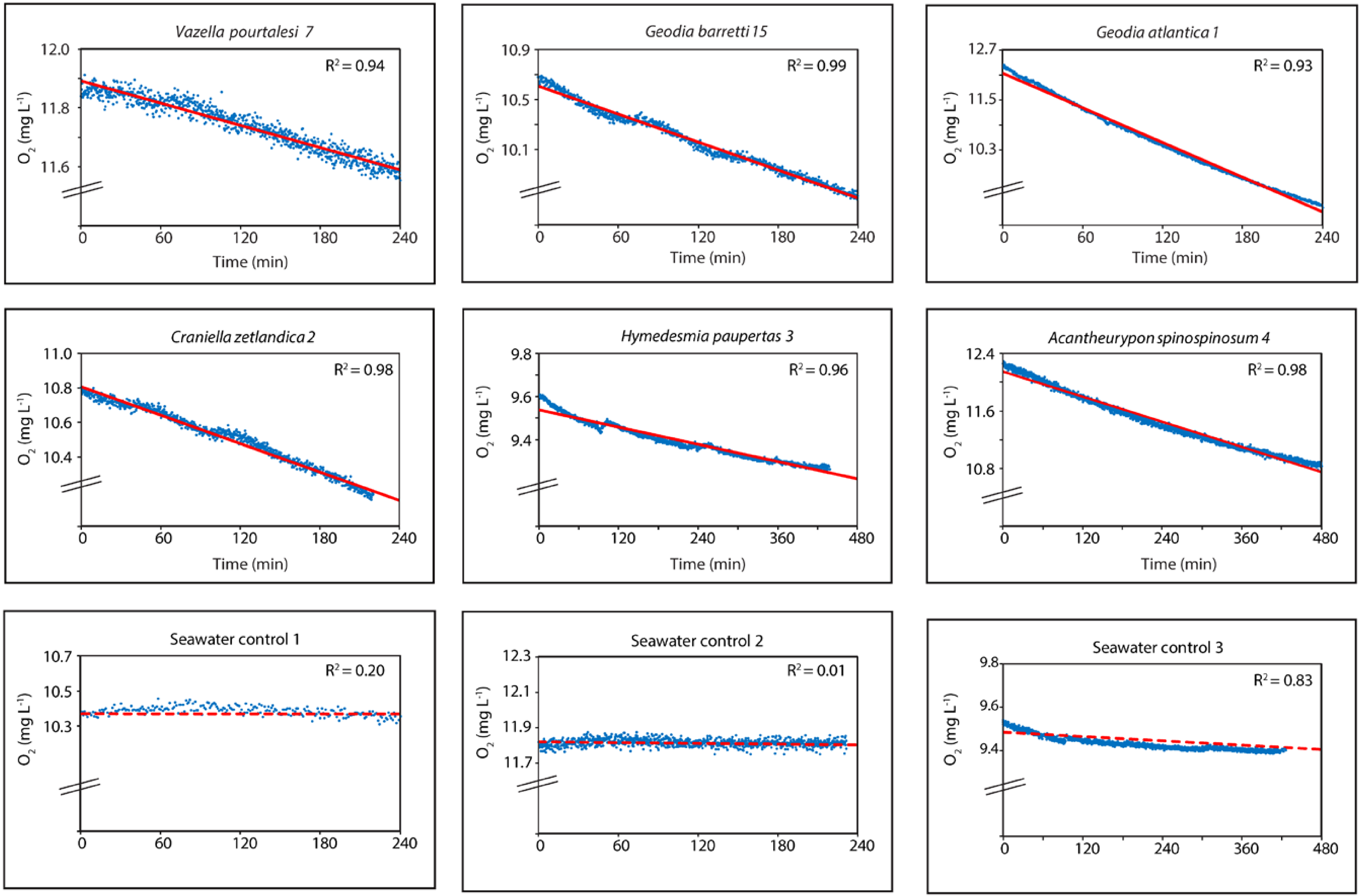

### Supplementary Figure S2

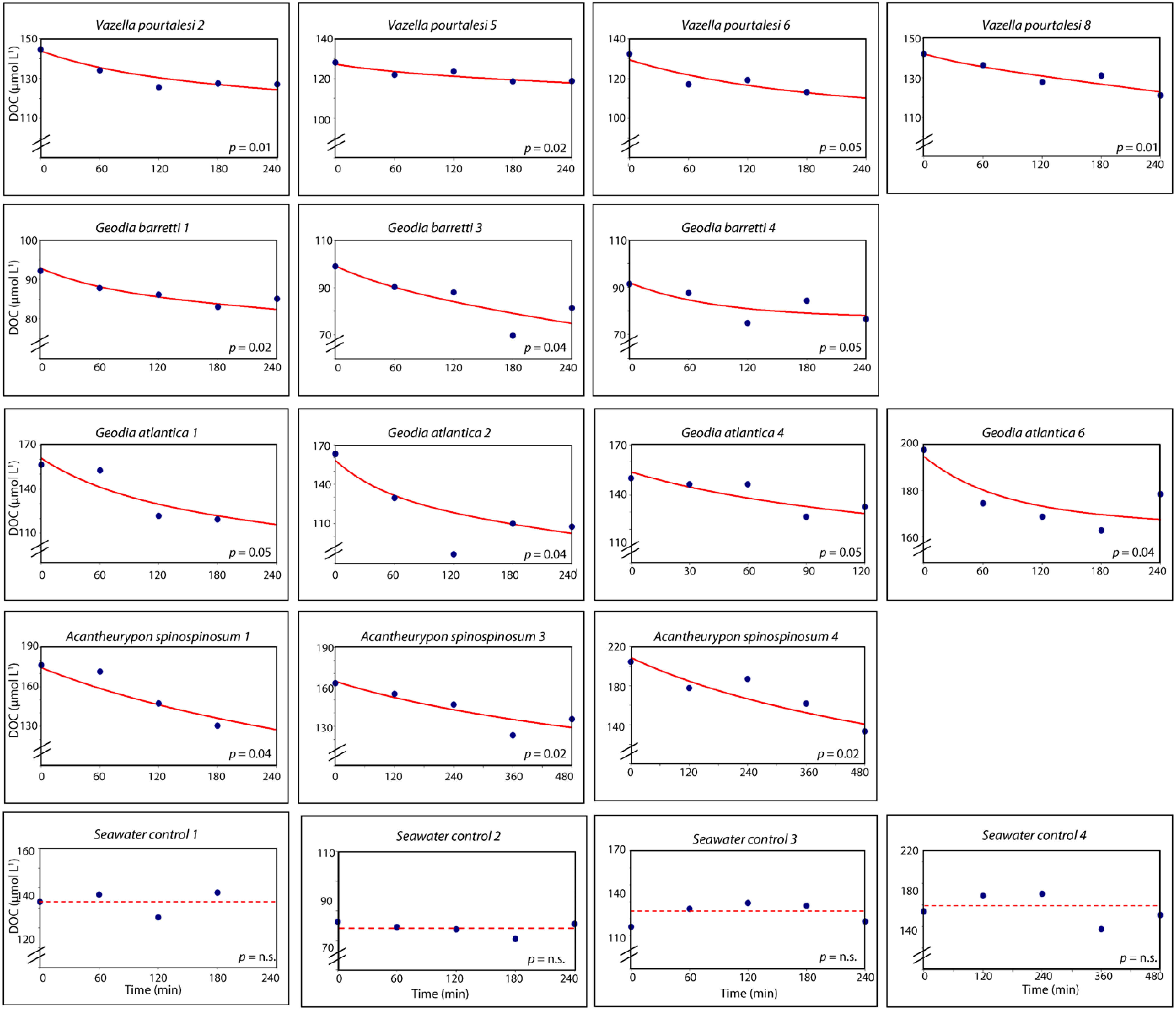
